# Compensatory and additive helper effects in the cooperatively breeding Seychelles warbler (*Acrocephalus sechellensis*)

**DOI:** 10.1101/372722

**Authors:** Lotte A. van Boheemen, Martijn Hammers, Sjouke A. Kingma, David S. Richardson, Terry Burke, Jan Komdeur, Hannah L. Dugdale

**Author notes:** Corresponding author: Lotte A. van Boheemen.

## Abstract

- In cooperatively breeding species, helper aid may affect dominant breeders’ investment trade-offs between current and future reproduction. By compensating for the care provided by helpers, breeders can reduce the costs of reproduction and improve chances of survival. Also, helper care can be additive to that of dominants, resulting in higher success of the current brood.
- However, the influence of helpers on offspring care itself may be the by-product of group size and territory quality. Therefore to make conclusive inferences about causation of additive and compensatory care as a result of help *per se* requires disentangling the impact of helping from other factors determining parental investment.
- In this study, we use 20 years of offspring provisioning data to investigate the effect of helping on breeder and overall offspring provisioning rates in the facultative cooperatively breeding Seychelles warbler (*Acrocephalus sechellensis*). Our extensive dataset allowed us to effectively control for the effects of living in a larger group and in territories with higher food availability.
- We show compensatory and additive care in response to helper aid. Helpers lightened the provisioning load of the dominant male and female and increased the total provisioning to the nestlings. This was irrespective of group size or territory quality (food availability).
- Our results illustrate how multiple benefits of helping behaviour can simultaneously be fundamental to the evolutionary maintenance of cooperative behaviour.

## Introduction

In cooperative breeding systems, offspring care is often shared between a dominant male and female ‘breeder’, and a variable number of subordinate helpers (Stacey & Koenig 1990; Solomon & French 1997; Koenig & Dickinson 2016; Komdeur *et al.* 2017). The optimal amount of parental investment provided by a breeder is determined by the trade-off between current and future reproduction (Williams 1966; Stearns 1989; Stearns 1992) and the care provided by helpers may affect the balance of this trade-off for the dominants. For example, care provided by helpers may increase the success of the current reproductive attempt, allow the dominants to reproduce more frequently, and/or improve the survival and future reproductive output of the dominants (Brown *et al.* 1978; Heinsohn 2004; Kingma *et al.* 2010; Koenig & Dickinson 2016).

The investment strategies implemented by cooperative breeders are generally classified as ‘additive’ and ‘compensatory’ care strategies (Hatchwell 1999). When the helpers improve overall care, the care provided by helpers is additive (Emlen & Wrege 1991). The resulting increase of the total amount of care received by the offspring can lead to higher reproductive success (Emlen & Wrege 1991; Komdeur 1994; Hatchwell 2004), through accelerated nestling growth (Dickinson *et al.* 1996) and reduced nestling starvation (Heinsohn 1995; Dickinson *et al.* 1996; Hatchwell 1999; Hatchwell 2004; Kingma *et al.* 2010). Conversely, when the dominants compensate for the care provided by helpers and reduce their amount of care, the total amount of care received by the offspring may remain similar. Such ‘load-lightening’ by helpers can reduce the costs of reproduction for the dominants (Heinsohn 2004; Meade *et al.* 2010; Koenig & Walters 2011), which can lead to increased dominant survival (Heinsohn 1992; Hatchwell & Russell 1996; Khan & Walters 2002; Cockburn *et al.* 2008; Kingma *et al.* 2010) and increased future reproductive success (Brown & Brown 1981; Woxfold & Magrath 2005; Blackmore & Heinsohn 2007; but see Meade *et al.* 2010).

These additive and compensatory investment strategies are not mutually exclusive and may occur at the same time (Hatchwell 1999; Kingma *et al.* 2010). For example, the dominants may reduce their offspring care investment in response to receiving help (load-lightening), but when this reduction is lower than the contribution of the helper the total amount of offspring care will increase (additive care). The relative importance of each of these strategies may be driven by the likelihood of offspring starvation, with more additive care when the risk of offspring starvation is higher, and more compensatory care when the risk of starvation is lower (Hatchwell 1999; Johnstone 2011).

Load-lightening and additive care strategies have been studied in many cooperative breeding systems (see Hatchwell 1999; Heinsohn 2004), but it is often extremely difficult to disentangle the effect of helpers from the effects of living in a larger group and from territory quality (Cockburn *et al.* 2008; Kingma *et al.* 2014). For example, larger groups with more helpers may be better able to occupy territories with higher food availability; hence the care to offspring might increase as a consequence of higher food availability in territories with helpers and not because of the effect of helpers *per se*. Similarly, load lightening of breeders could be the consequence of food being more difficult to find if more individuals occupy the territory and utilise the food, and in such cases breeders would not actually reduce the energy expenditure of providing care. Therefore, studies on load-lightening and additive care should attempt to disentangle the impact of helping from that of living in a larger group or in a territory with higher food availability.

Here, we use 20 years of parental and group provisioning data to investigate how helpers affect breeder and overall offspring provisioning rates in the facultative cooperatively breeding Seychelles warbler (*Acrocephalus sechellensis*). Seychelles warblers live in groups that occupy stable territories that are defended year-round (Komdeur 1991). Groups consist of a pair-bonded dominant male and female and 0-5 subordinate individuals of either sex that may or may not provide help with provisioning nestlings and fledglings (Komdeur 1994; Kingma *et al.* 2016). The presence of subordinate helpers and non-helping subordinates provides the opportunity to disentangle the impact of helping and group size (Cockburn *et al.* 2008). Subordinates are generally retained offspring from previous reproductive attempts in the territory (but see Richardson *et al.* 2007; Groenewoud *et al.* 2018). Dominant individuals gain from helper care as this positively influences first-year survival of offspring (Komdeur 1994), an effect that persists into adulthood (Brouwer *et al.* 2012). A previous study on a dataset collected during the first few years of the Seychelles warbler study found that (1) nests with helpers received a higher amount of total care compared to nests without helpers; (2) provisioning effort of dominant females was independent of helper presence; and (3) dominant males reduced their provisioning rates in groups with more helpers (Komdeur 1994). Here, we attempt to replicate this study, and, specifically, distinguish the impact of help from the effects of group size (including helpers and non-helpers) and food availability, using a much larger dataset collected during a subsequent time period (excluding the earlier years analysed in the previous study). Our results show that helpers provide both load-lightening and additive benefits to both dominant male and female Seychelles warblers.

## Methods

### Study population

The Seychelles warbler population on Cousin Island (29 ha; 04°20’ S, 55°40’ E) has been monitored closely since the mid-1980s (Komdeur *et al.* 2016). The main breeding season is July–September, and a smaller breeding season occurs January–March (Komdeur 1996). From 1997 onwards, ca. 96% of the population has been colour-ringed, using a unique combination of a metal British Trust for Ornithology ring and colour rings (Richardson *et al.* 2001). The identity of all colour-ringed birds present in each territory were recorded and the sex of all birds blood sampled since 1995 has been molecularly determined (Griffiths *et al.* 1998). Dominant birds, defined as the pair-bonded male and female in a territory based on their behavioural interactions (Richardson *et al.* 2002), form long-term pair bonds. Groups may contain 0-5 sexually mature (>5 months old) subordinates, which are usually retained offspring (Kingma *et al.* 2016; Groenewoud *et al.* 2018) and typically produce one clutch per season of a single-egg (87%; range 1–3 eggs) and nestlings fledge 18–20 days after hatching (Komdeur 1991). Subordinate birds were defined as ‘helpers’ when they were observed brooding or provisioning offspring. Territories were checked for breeding activity at least once every two weeks by following the dominant female for 15 minutes. Once breeding, focal territories were checked every week for at least 15 minutes to confirm for nest building, brooding or feeding activity.

### Provisioning observations

We measured nestling and fledgling provisioning rates at nests produced between 1995 and 2015. Provisioning watches with >10% of provisioning events by unidentified birds were excluded from the analyses (*N* = 178 of 701 watches). A total of 567 hours of 60–90 minute observations were included in our analyses (*N* = 523 watches). Provisioning rates were calculated as the number of nest visits during which the nestlings was fed. To account for variable types of observations, we grouped provisioning watches into three categories: *i*) provisioning and brooding: a nestling was fed in the nest and a female was still brooding; *ii*) provisioning nestling: a nestling was fed in the nest and no brooding occurred; and, *iii*) provisioning fledgling: a fledgling was fed away from the nest. Although brooding during provisioning can occur as a way to protect the nestling from the environment, most brooding occurred immediately after hatching (field observations).

### Monthly insect abundance and territory quality

Seychelles warblers are insectivorous, taking 98% of their insect food from the undersides of leaves (Komdeur & Pels 2005; Komdeur 2006). Insect abundance was estimated by counting the number of insects on the undersides of 50 leaves of the most abundant plant species (Komdeur 1991; Eikenaar *et al.* 2010), at 15 (until 1999) or 14 (after 1999) fixed locations on the island once every month. Monthly insect abundance was calculated as the mean insect abundance across these locations. The number of insects present in a territory is a useful index of territory quality (Komdeur 1994). For this, insect abundances in each territory were extrapolated from the nearest insect count location (Komdeur 1991). Leaf area was assessed by measuring the area of leaves of the abundant plant species (five leaves per species per site) at 50 random sites on the island (Komdeur 1991). Vegetation abundance was scored by determining the presence of all plant species at 20 random points in a territory in the following height bands: 0–0.75 m, 0.75–2 m, 2–4 m, and at 2-m intervals thereafter (Komdeur 1991). Territory sizes were measured each season using ArcGIS 9; territory boundaries were based on observations of individual warblers and the outcomes of territory disputes. Territory quality estimates were then calculated as insect abundance per unit leaf area (dm^2^) multiplied by vegetation abundance score, multiplied by territory size. These were then standardised across territories in each breeding season, by mean centering and dividing by two standard deviations (Gelman & Hill 2007). To provide an overall index of territory quality for each territory and investigate long-term effects of environment on investment, we calculated mean standardised territory quality per territory over all seasons (Hammers *et al.* 2012).

### Statistical methods

We performed generalized linear mixed model analyses in MCMCglmm 2.24 (Hadfield 2010), which takes a Bayesian approach, in R 3.4.0 (R Core Team 2017). We first investigated the impact of helper care on the dominants’ parental investment by modelling the number of provisioning visits by each dominant individual to offspring. Along with the number of helpers, we included the sex of the dominant individual, number of offspring, group size, provisioning watch type (provisioning and brooding, provisioning nestling, provisioning fledgling), monthly insect abundance and territory quality index as fixed effects. To explore sex differences in provisioning in response to helper presence or type of provisioning watch, we tested for an interaction between the number of helpers and sex of the dominant individual, and provisioning watch type and sex of the dominant individual. To account for varying observation duration, the log of the watch duration was also included in the fixed structure and a prior was specified to set its regression coefficient to 1 (i.e. observation duration was treated as an offset; J. Hadfield, personal communication). To control for repeated measures from dominant individuals that provisioned in more than one breeding season, we included bird identity as a random effect, using an idh variance structure to allow sex-specific variances to be estimated. To control for multiple provisioning watches and simultaneous watches of males and females at the same nest, we included the random effects of provisioning watch identity nested within nest identity. We did not include territory identity as most breeders live in the same territory during their entire life. To control for differences between observers we included observer identity as a random effect. For the random effects, we applied expanded priors (with V = 1, nu = 0.002, alpha.mu = 0, alpha.V = 1000) as the variance was close to zero (Hadfield 2015). For bird identity and residual variance, the expanded prior was structured as a 2×2 matrix to estimate variances for dominant males and females separately. The model had a Poisson error distribution and log link, was run for 4.5×10^5^ iterations with a burn-in of 5×10^4^ and thinning of 400. To test whether helper effects were additive or compensatory, we modelled the total number of provisioning visits per watch by all feeders (i.e. the dominants and helpers) to offspring. This model was the same as the provisioning model except that the response was the total number of feeds, the parameters describing sex and bird identity were omitted and the model ran for 2.1×10^7^ iterations with a 1×10^6^ burn-in and 2×10^3^ thinning. Provisioning observations of nests with more than one nestling can be confounded by factors such as sibling competition (Bebbington *et al.* 2017) and reduced statistical power resulting from low sample size of nests with more than one nestling (48/523). We therefore ran additional models with identical settings, excluding number of offspring as a fixed effect.

To assess model convergence, we checked that the: *i*) autocorrelation for all parameters was <0.1; *ii*) variance estimates passed the Heidelberger and Welch’s convergence diagnostic, which tests if successive samples are drawn from a stationary distribution; *iii*) variance estimates passed the Geweke diagnostic, which tests for equality of the means of the first 10% and last 50% of the Markov chain; and *iv*) variance inflation between fixed effects was <3 to avoid colinearity (Heidelberger & Welch 1983; Geweke 1991; Cowles & Carlin 1996). We evaluated if the 95% credibility intervals of the posterior modes overlapped zero, where a departure from zero was interpreted as a significant effect.

## Results

Both male and female dominants showed lower provisioning effort when more helpers aided in provisioning (12.6% reduction in predicted feeds/hour per helper, from 8.5 (no helpers, *N* = 492) to 8.1 (one helper; *N* = 350) and 7.4 feeds/hour (two helpers; *N* = 47); Fig. 1 & 2). This load-lightening effect was similar for males and females as the 95% CI’s of the interaction between the sex of the dominant and the number of helpers included zero (Fig. 1). An interaction between the sex of the dominant and provisioning watch type revealed that the provisioning rates of dominant males were 27.0% higher to nestlings (8.0 feeds/hour) versus fledglings (5.8 feeds/hour; Fig. 1 & 3). The opposite pattern was observed in dominant females, which fed fledglings almost twice as much as nestlings (12.0 versus 6.8 feeds/hour; Fig. 3). Feeding rates were not significantly related to monthly insect abundance, territory quality, number of offspring, or group size (Fig. 1).

**Figure 1.**
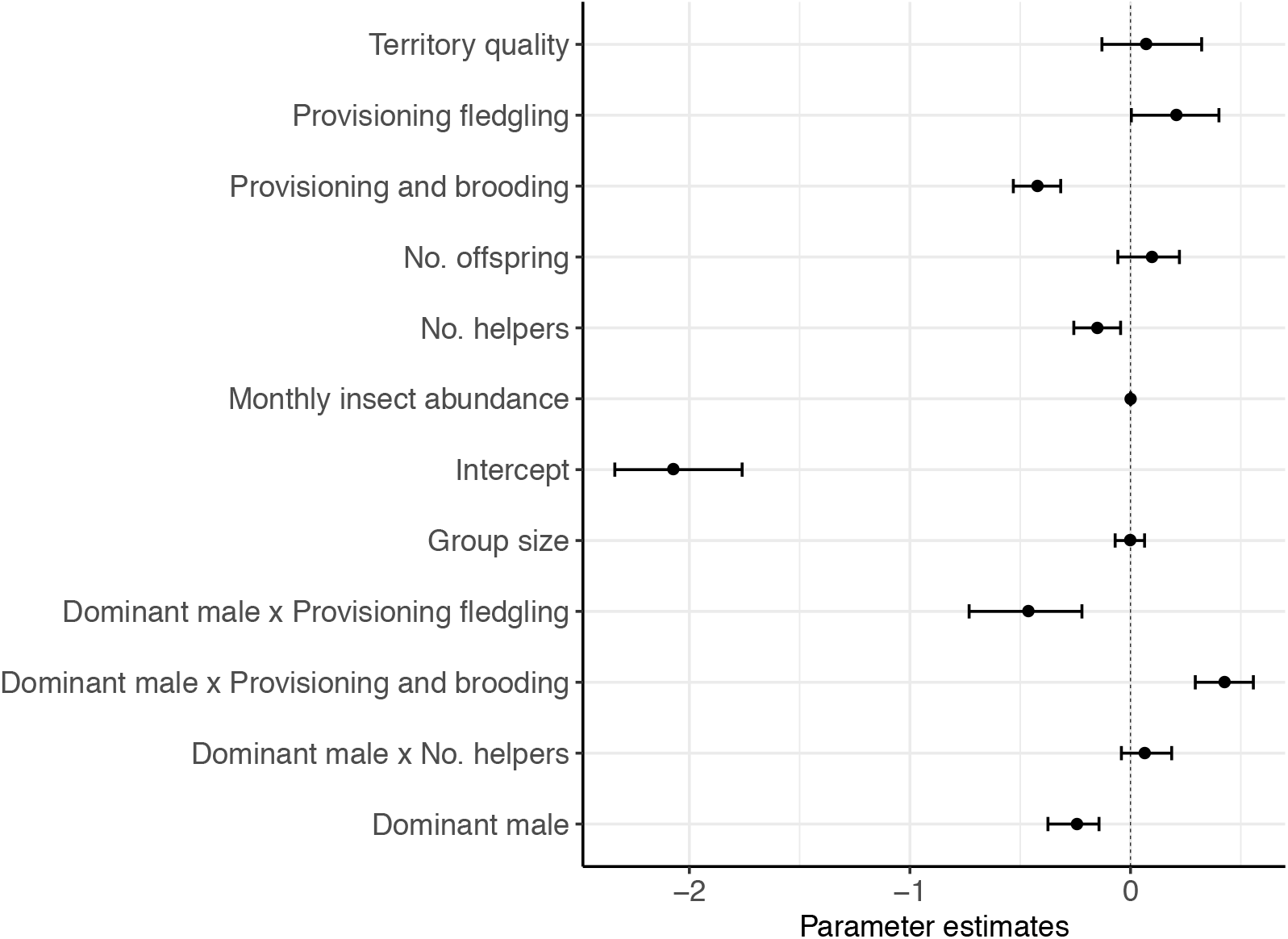
Posterior density estimates of parameter modes, and their 95% credible intervals, for the fixed effects used to model the number of feeds by dominant birds: watch type (provisioning and brooding = 438, provisioning nestling = 384, provisioning fledgling = 67; contrast = provisioning nestling), sex of the dominant bird (male = 446, female = 443; contrast = female), number of helpers (0 = 492, 1 = 350, 2 = 47), group size (2 = 280, 3 = 425, 4 = 150, 5 = 30, 6 = 4), number of offspring (1 = 808, 2 = 79, 3 = 2), index of territory quality monthly insect abundance, watch duration. * indicates parameters whose credible intervals do not overlap zero.

**Figure 2.**
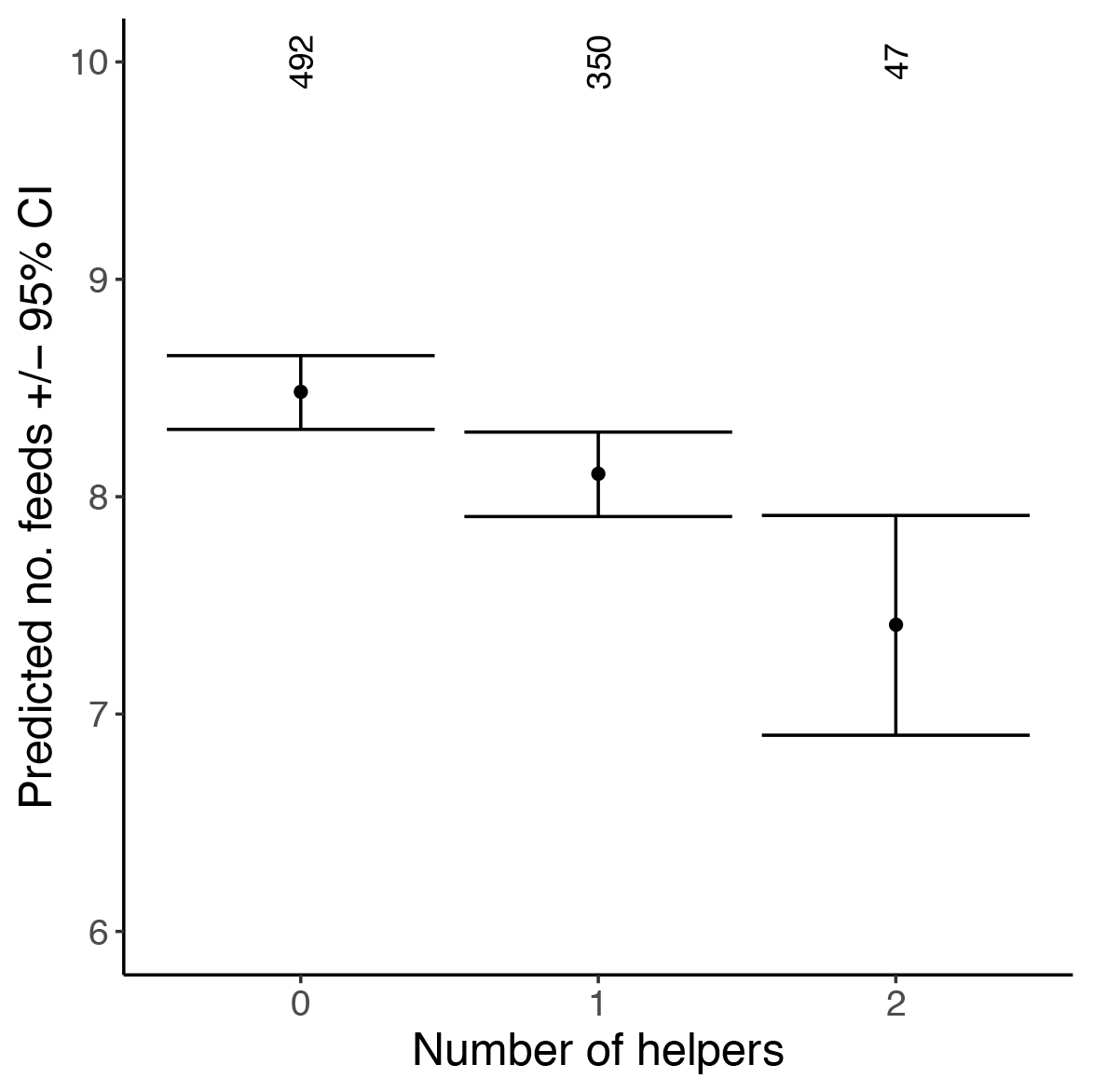
The mean number of feeds by dominant birds related to number of helpers present. Error bars represent 95% confidence intervals and numbers at the top of the graph represent sample sizes.

**Figure 3.**
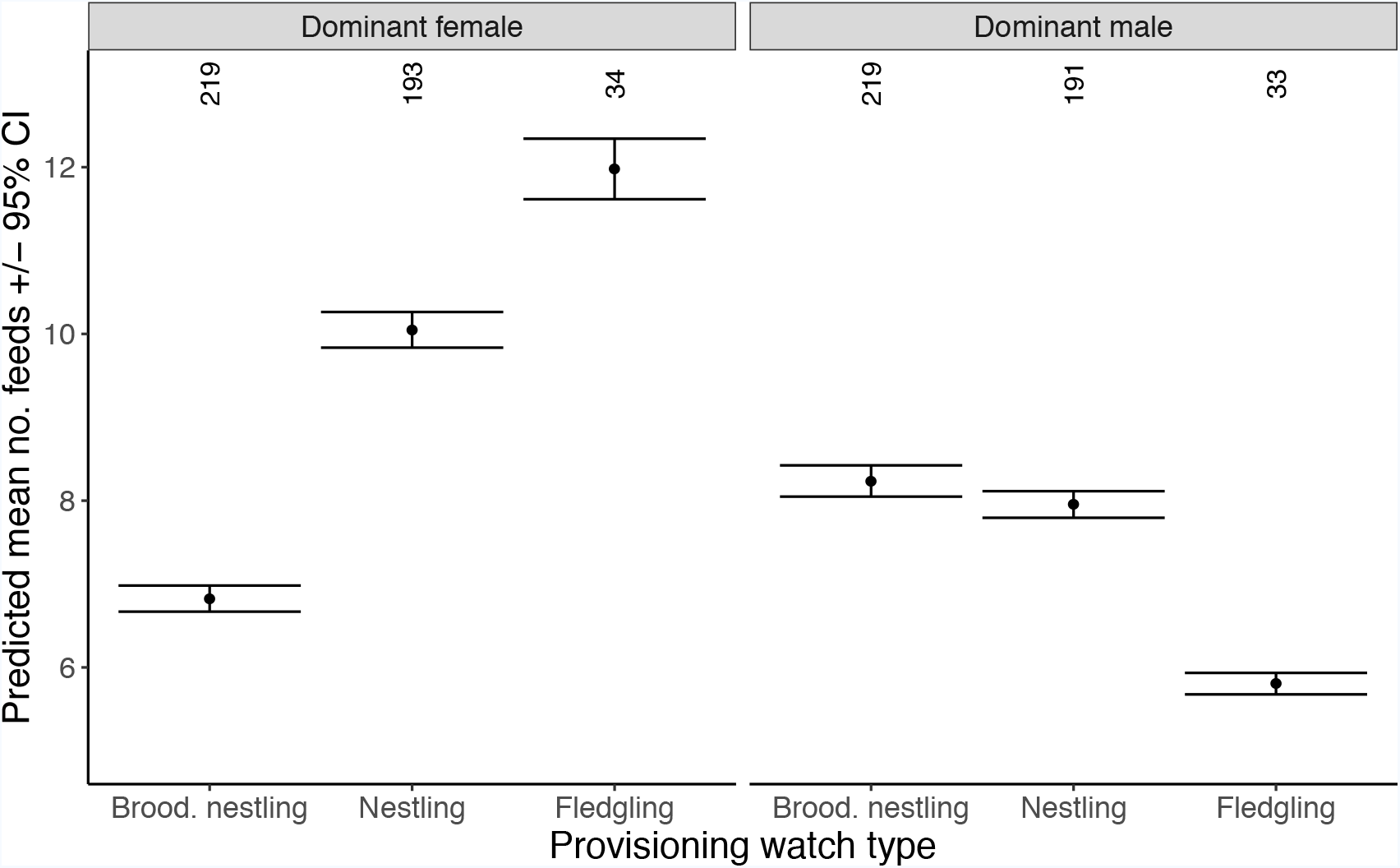
The mean number of feeds by dominant males and females in relation to the three types of provisioning watches (provisioning & brooding nestlings (brood. nestling), provisioning nestlings (nestling) and provisioning fledglings (fledgling)). Error bars represent 95% confidence intervals.

We found a strong increase in total provisioning when helpers were feeding and also when more helpers were involved (Fig. 4). A single helper resulted in a predicted increase of 30.5% (22.2 visits per hour, *N* = 177, compared to 17.0 feeds in pairs, *N* = 248) provisioning visits per hour, and a second helper increased the total provisioning effect to 64.7% increase (28.0 feed/hour, *N* = 24; Fig. 5). The total number of provisioning visits each hour to nestlings also being brooded was 23.0% less than to nestlings only being provisioned (17.6 versus 21.6; Fig. 4 & 6). The total number of provisioning visits received by offspring was not correlated with group size, number of offspring, territory quality or monthly insect abundance (Fig. 4). Excluding number of offspring from these models did not change the direction or significance of our results (Supporting information). Together, these results indicate load-lightening and total provisioning increased with additive feeding investment by helpers.

**Figure 4.**
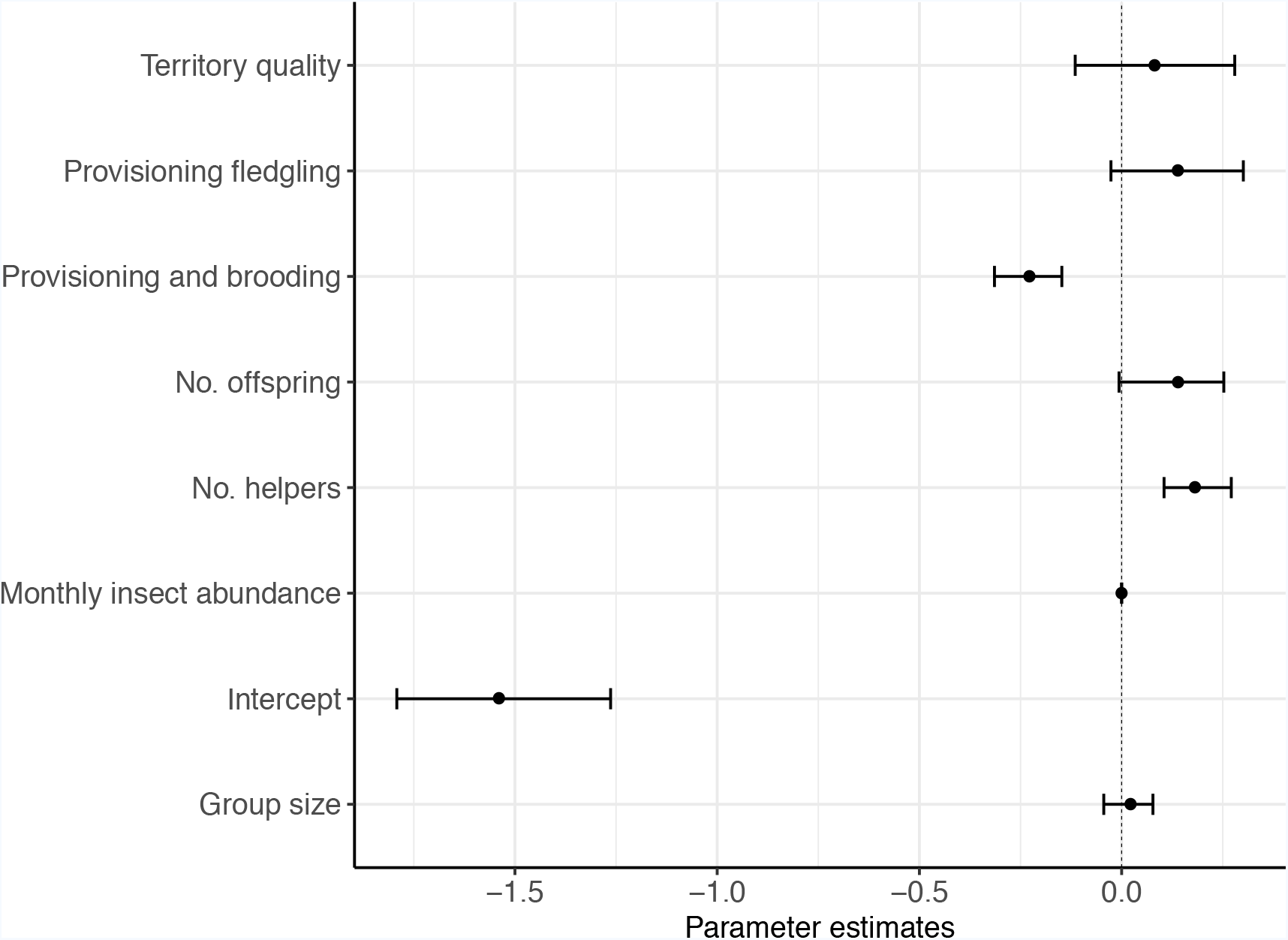
Posterior density estimates of parameter modes, and their 95% credible intervals, for the fixed effects used to model the total number of feeds per provisioning watch: watch type (provisioning and brooding = 222, provisioning nestling = 193, provisioning fledgling = 34; contrast = provisioning nestling), number of helpers (0 = 248, 1 = 177, 2 = 24), group size (2 = 141, 3 = 214, 4 = 77, 5 = 15, 6 = 3), number of offspring (1 = 408, 2 = 40, 3 = 1), index of territory quality, monthly insect abundance, watch duration. * indicates parameters whose credible intervals do not overlap zero.

**Figure 5.**
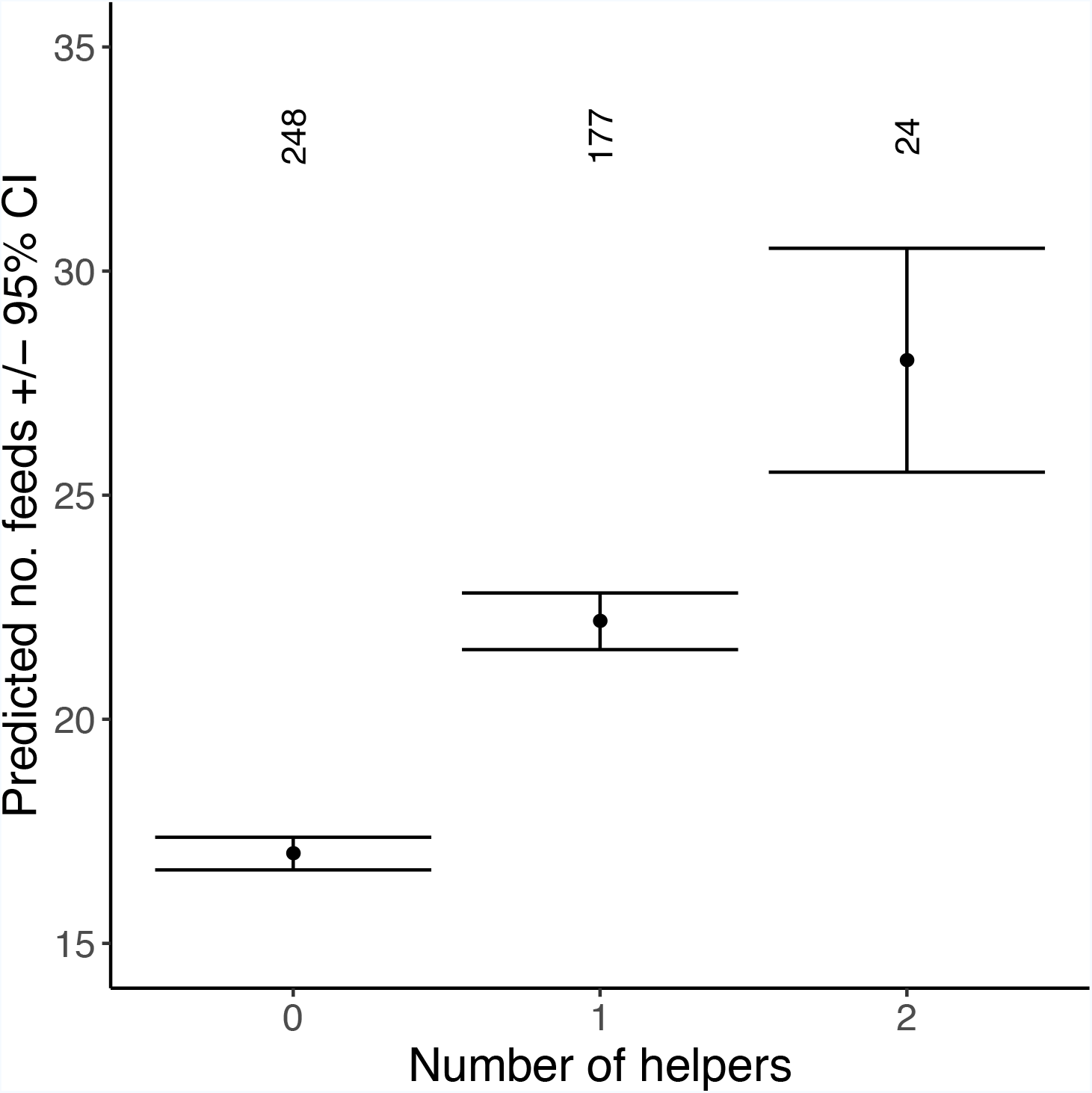
The total number of feeds to the nest in relation to the number of helpers present. Error bars represent 95% confidence intervals. Numbers at the top of the graph represent sample sizes.

**Figure 6.**
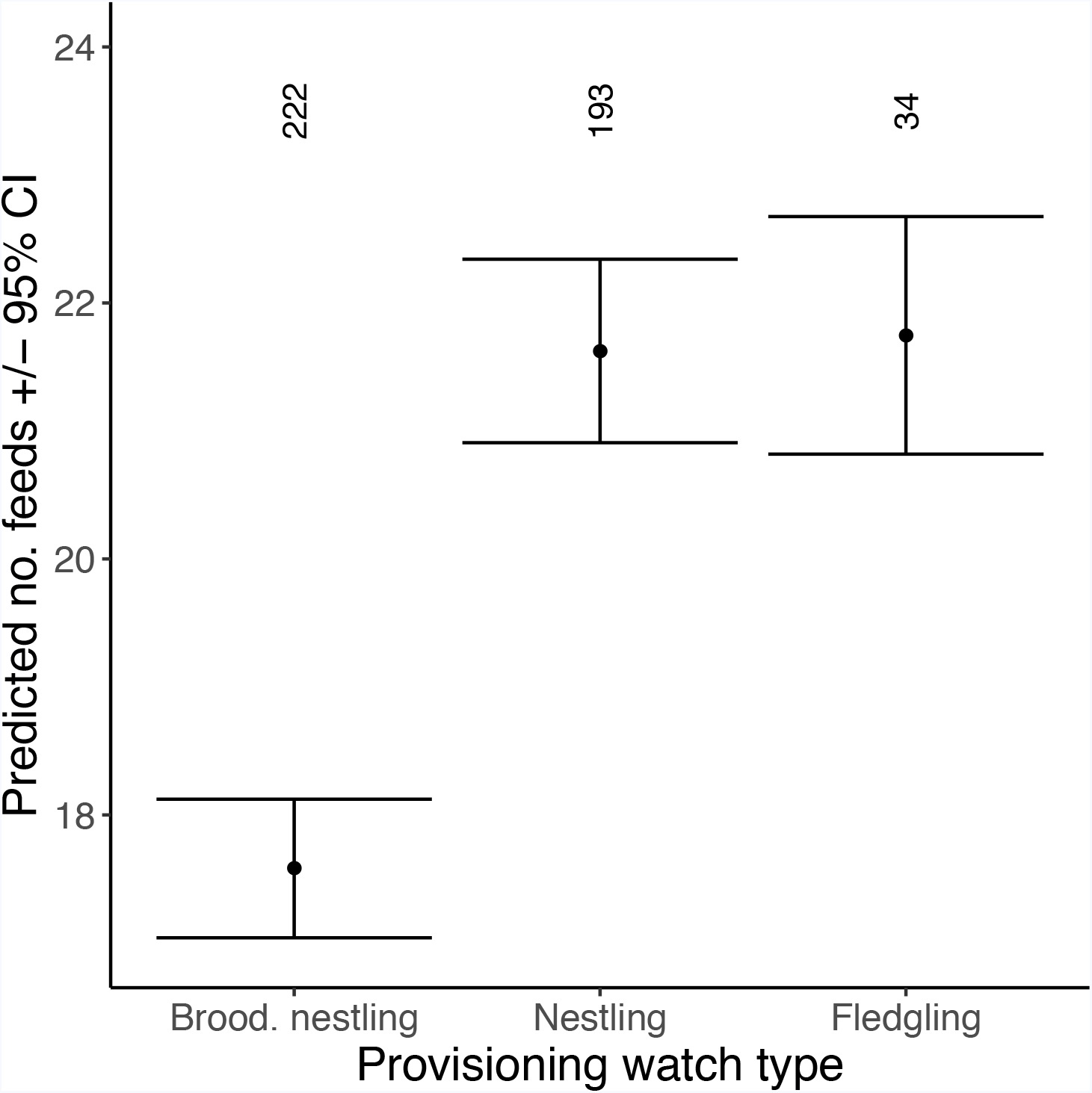
The total number of feeds received by the offspring in relation to the three types of provisioning watches (provisioning & brooding nestlings (brood. nestling), provisioning nestlings (nestling) and provisioning fledglings (fledgling)). Error bars represent 95% confidence intervals. Numbers at the top of the graph represent sample sizes.

## DISCUSSION

Our analysis of the long-term Seychelles warbler data set revealed both additive and compensatory helper effects in this species. Helpers lightened the provisioning load of dominant individuals and increased the total number of provisioning trips to the nestlings. These results were not the spurious result of group size or territory quality. Moreover, as non-dominants could be defined as helpers when aiding in brooding only, this is a conservative analysis and the actual additive and compensatory effects might be higher. The increased total nest provisioning effort resulting from additive helper provisioning could lead to higher nestling survival (Hatchwell 1999; MacColl & Hatchwell 2003; Woxfold & Magrath 2005; Valencia *et al.* 2006). Indeed, in the Seychelles warbler, this may well explain the higher survival of offspring in their first year (Komdeur 1994) and beyond (Brouwer *et al.* 2012) leading to direct fitness benefits for parents.

We demonstrated that, in addition to additive care, helpers also provide load-lightening benefits for dominant individuals, as dominants of both sexes reduced provisioning rates when aided by helpers. In some, but not all, species (Heinsohn 2004; Kingma *et al.* 2010) such load-lightening benefits have been associated with increased survival of dominants with helpers. In the Seychelles warbler, survival of dominants with and without helpers is similar (Komdeur 1994; Hammers et al. in revision), except among very old dominants when those that receive help show higher survival (Hammers et al. in revision). While it may thus be that load-lightening effects on breeder survival are only obvious in some circumstances (i.e. when breeders are old), other reproductive components (like re-nesting opportunities or time between nesting attempts) may also be affected by breeders reducing their current workload. Future work will need to reveal whether such effects may explain selection on breeders reducing workload in response to help.

We found that, overall, the provisioning rates of male dominants were lower than those of female dominants and these sex-specific provisioning rates varied with the type of provisioning watch. Sex-related differences in parental investment of the dominants are not uncommon (Hatchwell 1999; MacColl & Hatchwell 2003) and are proposed to result from diverging cost-benefits trade-offs between the sexes (MacColl & Hatchwell 2003). Several studies have shown that the genetic relatedness of the carer to the brood affects investment, where male uncertainty of parentage can result in lower amounts of care (e.g.Burke *et al.* 1989; Neff 2003; Kokko & Jennions 2012). In the Seychelles warbler, male breeders are on average less related to the offspring than females, due to the 44% extra-pair paternity occurring in this species (Richardson *et al.* 2001; Hadfield *et al.* 2006), which may explain the overall lower provisioning by breeder males. The frequency of male extra-pair copulations might change over the course of the breeding event, with possible higher opportunity during the fledgling state, reflecting their lower provisioning effort at this time.

The observation that sex-specific investment changed over the course of the breeding event may suggest that other aspects, besides certainty of parentage, affect the asymmetry in provisioning between sexes, as has been observed in other species (Cockburn *et al.* 2008; Meade *et al.* 2010). For example, females might reduce the costs of investment before the nestling period by decreasing egg size when assisted by helpers (Russell *et al.* 2008; Dixit *et al.* 2017; but see Koenig *et al.* 2009). In the Seychelles warbler, females predominantly build the nest and brood the egg, and spend less time foraging compared to males, who guard the nest (Komdeur & Kats 1999). This, in combination with on-going brooding of newly hatched chicks, may suggest higher costs for females during the pre-nestling and young-nestling period, which could explain lower provisioning effort of the dominant female compared to the dominant male shortly after hatching. The most suitable investment strategy is therefore suggested to change within the breeding season and fine-scaled studies are required to understand the evolution of parental care (Savage *et al.* 2017).

Our results differ from previous findings of provisioning effort in the Seychelles warbler in relation to helper presence. Komdeur (1994)found a load-lightening effect for dominant males only when three or more helpers were present. The relatively higher degree of load-lightening identified here, for both sexes and with a smaller number of helpers, suggests that the cost-benefit trade-offs for dominant individuals may have changed since Komdeur’s earlier Seychelles warbler study. For instance, an increase in offspring survival (e.g. due to higher quality of insects or increased protection from the environment; Komdeur & Pels 2005) would allow parents to relax investment into the current brood. The emergence of such temporal variation in parental care raises further questions about the evolution of cooperative breeding.

## Conclusion

Altogether, our study adds to the growing evidence that both compensatory and additive care can apply at the same time within one species. These simultaneous parental care strategies are fundamental to the evolutionary maintenance of cooperative behaviour. The exact fitness effects of both load-lightening and additive care, as well as sex-specific changes in fitness benefits during the breeding season need to be explored in future.

## Acknowledgements

Nature Seychelles kindly allowed us to work on Cousin Island and provided accommodation and facilities during our stay. The Department of Environment and the Seychelles Bureau of Standards gave permission for fieldwork and sampling. We would like to thank the numerous researchers, students and field assistants for their invaluable help with data collection. HLD was funded by a NERC post-doctoral fellowship (NE/I021748/1), and LAB received funding from the Groningen University Fund and Marco Polo Fund, SAK and MH by a Veni fellowship by NWO.

